# Deep Learning Global Glomerulosclerosis in Transplant Kidney Frozen Sections

**DOI:** 10.1101/292789

**Authors:** Jon N. Marsh, Matthew K. Matlock, Satoru Kudose, Ta-Chiang Liu, Thaddeus S. Stappenbeck, Joseph P. Gaut, S. Joshua Swamidass

**Affiliations:** J. N. Marsh and S. J. Swamidass are with the Department of Pathol-ogy & Immunology and the Institute for Informatics at Washington University School of Medicine, St. Louis, MO, 63110 USA e-mail: (see http://swami.wustl.edu/contact). *(Jon N. Marsh and Matthew K. Matlock are co-first authors.)*; M. K. Mattlock, S. Kudose, T.-C. Liu, T. Stappenbeck, and J. Gaut are with the Department of Pathology & Immunology at Washington University in St. Louis, St. Louis, MO, 63110 USA.

**Keywords:** kidney, glomerulosclerosis, digital pathology, fully convolutional network, donor organ evaluation

## Abstract

Transplantable kidneys are in very limited supply. Accurate viability assessment prior to transplantation could minimize organ discard. Rapid and accurate evaluation of intra-operative donor kidney biopsies is essential for determining which kidneys are eligible for transplantation. The criteria for accepting or rejecting donor kidneys relies heavily on pathologist determination of the percent of glomeruli (determined from a frozen section) that are normal and sclerotic. This percentage is a critical measurement that correlates with transplant outcome. Inter- and intra-observer variability in donor biopsy evaluation is, however, significant. An automated method for determination of percent global glomerulosclerosis could prove useful in decreasing evaluation variability, increasing throughput, and easing the burden on pathologists. Here, we describe the development of a deep learning model that identifies and classifies non-sclerosed and sclerosed glomeruli in whole-slide images of donor kidney frozen section biopsies. This model extends a convolutional neural network (CNN) pre-trained on a large database of digital images. The extended model, when trained on just 48 whole slide images, exhibits slide-level evaluation performance on par with expert renal pathologists. The model substantially outperforms a model trained on image patches of isolated glomeruli. Encouragingly, the model’s performance is robust to slide preparation artifacts associated with frozen section preparation. As the first model reported that identifies and classifies normal and sclerotic glomeruli in frozen kidney sections, and thus the first model reported in the literature relevant to kidney transplantation, it may become an essential part of donor kidney biopsy evaluation in the clinical setting.

## I. Introduction

HERE is a global shortage of donor kidneys suitable for transplantation exacerbated by an unacceptably high discard rate of recovered organs. Intra-operative examination of donor kidney biopsy frozen sections is essential to assess organ viability prior to transplantation. Many evaluation metrics are utilized, including percent global glomerulosclerosis, interstitial fibrosis, arteriosclerosis, and arteriolar hyalinosis [1]. The increased use of “expanded criteria donors” who are older and/or have comorbidities renders accurate evaluation of these pathologic findings increasingly important [1]–[4]. However, variability in biopsy evaluation between observers and institutions is distressingly large [1], [5]–[7], which may explain why these histologic features do not consistently correlate with outcome [1], [7]. Such variability may be heightened in the time-sensitive context of daily practice, where biopsies are often read by non-specialist pathologists at odd hours using frozen sections. Poor reproducibility amongst pathologists minimizes the utility of intraoperative organ assessment and may contribute to unnecessary organ discard. There is thus a need for new objective techniques to assist pathologists with rapid intraoperative donor kidney biopsy interpretation.

Identification of non-sclerotic and sclerotic glomeruli is an essential task that is required to compute percent global glomerulosclerosis, a critical feature that correlates with graft outcome [3], [7]–[9]. The United Network for Organ Sharing (UNOS) guidelines emphasize percent global glomerulosclerosis as a key factor in determining organ acceptance. While a variety of approaches have been described for automatic identification of glomeruli, none to our knowledge have addressed the challenges associated with intraoperative biopsy assessment for organ transplant. Logistical and time constraints in this setting often necessitate the use of frozen sections and their concomitant artifacts (e.g. cracking, holes), and H&E stains that are not optimized for visual differentiation of glomeruli from interstitial tissue (e.g. Figure 1 and Figure 2). These difficulties are compounded by the sheer volume of data present in gigapixel whole-slide images, which necessitate highly optimized algorithms to yield results in a timely fashion.

**Fig. 1:**
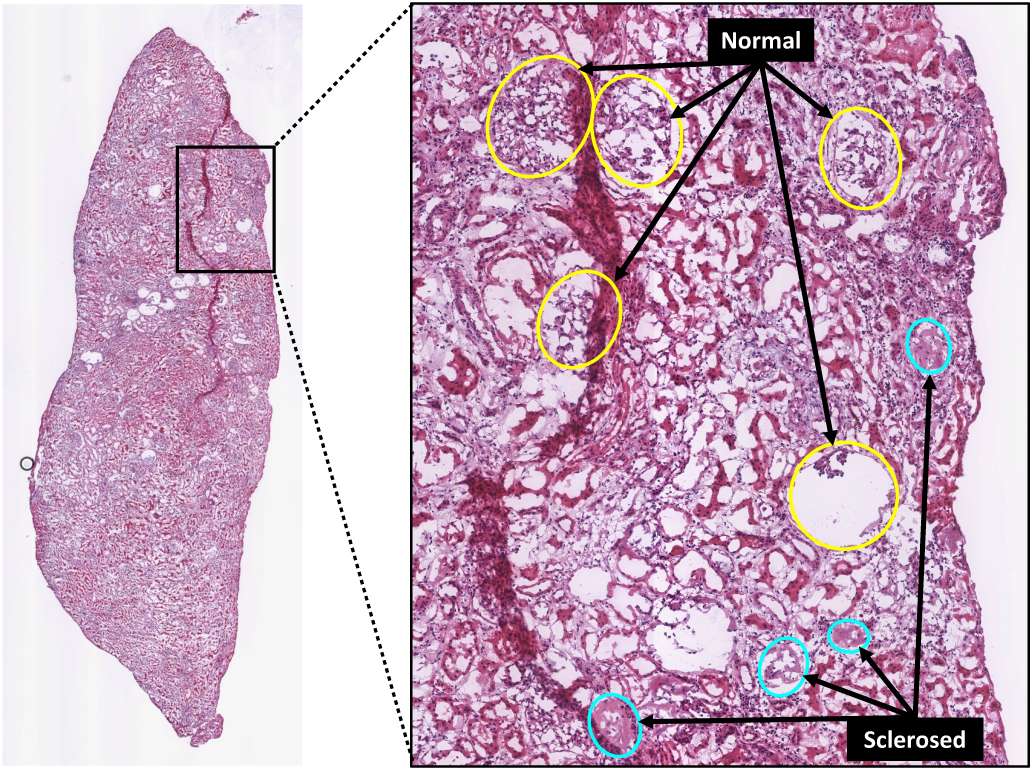
Example whole-slide image (WSI) of H&E-stained human renal frozen wedge biopsy scanned at 20X, with inset showing normal (yellow) and sclerosed (cyan) glomeruli as labeled by trained observers. Note the variability of appearance of glomeruli between and within categories.

**Fig. 2:**
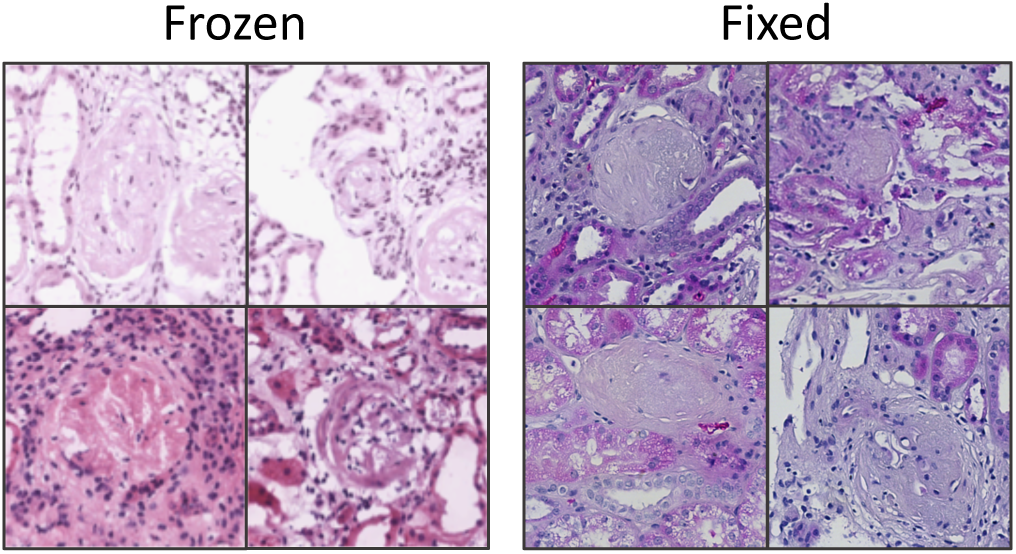
Example image patches (at 20X magnification) of globally sclerosed glomeruli from frozen (left) and formalin-fixed (right) H&E slide preparations. Variability in glomerular appearance and stain intensity is greater in frozen preparations. Also note variability in stain intensity in frozen samples, typical of the dataset used in this study.

The remarkable success of convolutional neural nets (CNNs) in generalized image recognition tasks suggests pathways to solving this seemingly intractable problem [10]–[13]. CNNs’ primary advantage is that the models automatically learn salient features from the data alone, rather than requiring a set of handcrafted parameters and extensive input normalization. The increasingly widespread use of CNNs has been facilitated by the concept of transfer learning, in which deep learning models, previously trained to categorize or identify objects in images from one domain, are repurposed for application in another. This is typically accomplished by freezing most, if not all, of an image-recognition network’s learned weights (which presumably encode a large number of generalized image features) below the classification layer, and then training the remaining layers to recognize features specific to the new domain. This leverages the vast amount of computational resources needed to train the model from scratch using randomly initialized weights on millions of input images; furthermore, fewer training examples are typically required for the repurposed model to converge on an optimized set of model weights, and training time can be significantly shortened. Often, investigators have adapted one of several CNNs trained on the ImageNet database [14] as the basis for medical image recognition algorithms. In the realm of histopathology, CNNs have been most commonly applied to cancer detection and classification [15]–[21]. More recently, several studies have employed CNNs for glomerulus identification in renal biopsies [22]–[25].

However, none of these studies describes the use of frozen sections as input to detection algorithms, nor are they capable of differentiating normal from sclerotic glomeruli after detection without the use of special stains. These studies, consequently, fall short of addressing the combination of issues associated with transplant evaluation.

In this study, we sought to evaluate the performance of CNN variants derived from a pre-trained image recognition network applied to the problem of glomerular identification and classification in renal preimplantation frozen section wedge biopsies. We compared a conventional patch-based CNN model with a fully convolutional CNN model. We show that the fully convolutional CNN model is superior and can be quickly trained on a relatively small dataset to yield results on par with expert renal pathologist interpretation.

## II. Related work

Detection of glomeruli in digitized histological images has been approached using a variety of methods. A summary of relevant details of recent work is shown in Table I.

**Table I.** Summary of related work (this study is represented in the top row). dashes represent instances where information was unreported.

| Ref | Imaging | Species | Task |  |
| --- | --- | --- | --- | --- |
| * | WSI, H&E, frozen | human | segmentation, classification | (normal, sclerosed) |
| [26] | WSI, anti-desmin, fixed | rat | segmentation, classification, characterization | (normal, diabetic) |
| [27] | WSI, anti-desmin, fixed | rat | segmentation | (normal, diabetic) |
| [28] | field, anti-nestin, fixed | rat | segmentation, characterization | (normal, renal failure) |
| [23] | WSI, PAS, fixed | mouse | segmentation | (normal) |
| [25] | WSI, Jones H&E/PAS/other, fixed | human | segmentation | (normal) |
| [22] | WSI, H&E/PAS/others | human, mouse, rat | segmentation, characterization | (normal, diabetic) |
| [29] | WSI, H&E | primate | segmentation, characterization | (normal, diseased) |
| [30] | WSI, Masson's trichrome | human | segmentation | - |
| [31] | field, H&E | mouse | segmentation | - |
| [24] | patch, PAS | human | segmentation | - |
| [32] | patch | rat | segmentation, characterization | (normal, hypertrophy) |
| [33] | patch, H&E/PAS, fixed | human | segmentation, characterization | (normal, proliferating) |
| [34], [35] | patch, H&E/PAS, fixed | mouse, rat | segmentation, characterization | (normal) |
| [36]–[38] | patch | - | segmentation | - |

The majority of studies incorporate domain-specific morphometric or texture-based techniques to search for and define glomerular boundaries. Many of these have demonstrated boundary detection for small image patches containing isolated glomeruli [31]–[33], [36]–[38]. Translating detection techniques to whole-slide images (WSI) containing numerous glomeruli is a necessary but more difficult undertaking. The task of detection over large image regions can be facilitated using immunohistochemical stains such as nestin [28] and desmin [26], [27] to highlight glomerular podocytes and enhance the utility of segmentation algorithms. However, the immunohistological approach is less applicable for evaluation of preimplantation biopsies. Other groups have used a variety of techniques for glomerulus identification on routine stains [10], [22], [29], [30], [34], [35], typically through the use of some combination of colorspace transformation, thresholding, and/or morphological descriptors alone or as input to support vector machines or CNNs.

Most recently CNNs have been explored as primary tools for glomeruli detection. Proof-of-concept classifiers adapted from both the AlexNet [39] and GoogleNet [10] models were shown to be able to differentiate image patches containing isolated normal glomeruli from non-glomerular structures [24]. CNNs were also demonstrated to outperform HOG classifiers in glomerulus detection accuracy when applied to random image patches from kidney WSI [25]. Additional promising results were demonstrated by cascading the output of one CNN (optimized for glomerulus detection in downsampled WSI) to another (adapted for precise segmentation at higher resolutions) [23]. This combination outperformed single CNNs in segmenting glomeruli; notably, the CNNs used a fully convolutional model based on the U-Net architecture [40] to yield pixel-mapped outputs, enabling “end-to-end” training on image patches randomly sampled over WSI.

It should be noted that none of these studies describes the use of frozen sections as input to detection algorithms. Likewise, in studies that characterized pathologic kidney samples [22], [26]–[29], [32], [33], only one method differentiated normal from pathologic glomeruli after detection, and this required the use of specialized immunostaining that highlighted damaged glomeruli [26]. The remainder describe differences in descriptors in glomeruli from normal and pathologic populations, rather than isolating and labeling different glomeruli within the same wide image field. The current study demonstrates the novel use of CNNs applied to frozen H&E sections to detect non-sclerotic and sclerotic glomeruli to assist pathologists in intra-operative interpretation of percent global glomerulosclerosis.

## III. Methods

### A. Data

WSIs were acquired from H&E-stained frozen wedge donor biopsies retrieved between April 2015 and July 2017 using the Washington University Digital Pathology Exchange (WUPAX) laboratory information system. Sections were scanned at 20x using an Aperio Scanscope CS scanner and stored in SVS format, then converted to TIFF format at full resolution (0.495 *µ*m/pixel). 48 sample WSIs (ranging in size from 187 megapixels to 482 megapixels) were acquired from the database and selected so as to exhibit a wide range of values of percentage globally sclerosed glomeruli (1% to 72%). The WSIs were obtained from a total of 20 kidneys recovered from 17 donors. The average total number of glomeruli was 81 ± 31 per WSI. Annotations used for training and testing the CNNs were initially created by a senior resident (SK) and subsequently amended by a board-certified renal pathologist (JPG). Annotation was performed manually by outlining and labeling all glomeruli (using elliptically shaped masks) in each WSI using an in-house plugin written for Fiji [41], in order to generate pixel-wise label masks of glomerulus regions at the same resolution as the parent WSI. All glomeruli were categorized into globally sclerotic (defined as sclerosis involving the entire glomerular tuft) or non-sclerotic (defined as any glomeruli that did not show global sclerosis). The globally sclerotic category included all types of global sclerosis: obsolescent, solidified and disappearing [42]. All non-glomerular areas (including tubules, vessels, inflammatory cells, and interstitium) were labeled as tubulo-interstitium. A total of 870 sclerosed and 2997 non-sclerosed glomeruli were labeled. Model training and testing was performed by grouping slides and their associated data into training and validation sets in a 6-fold cross-validation scheme. No image preprocessing was performed prior to training or testing.

### B. Models

Two models and training methodologies were used for glomeruli detection in WSI (see Figure 3):

**Fig. 3:**
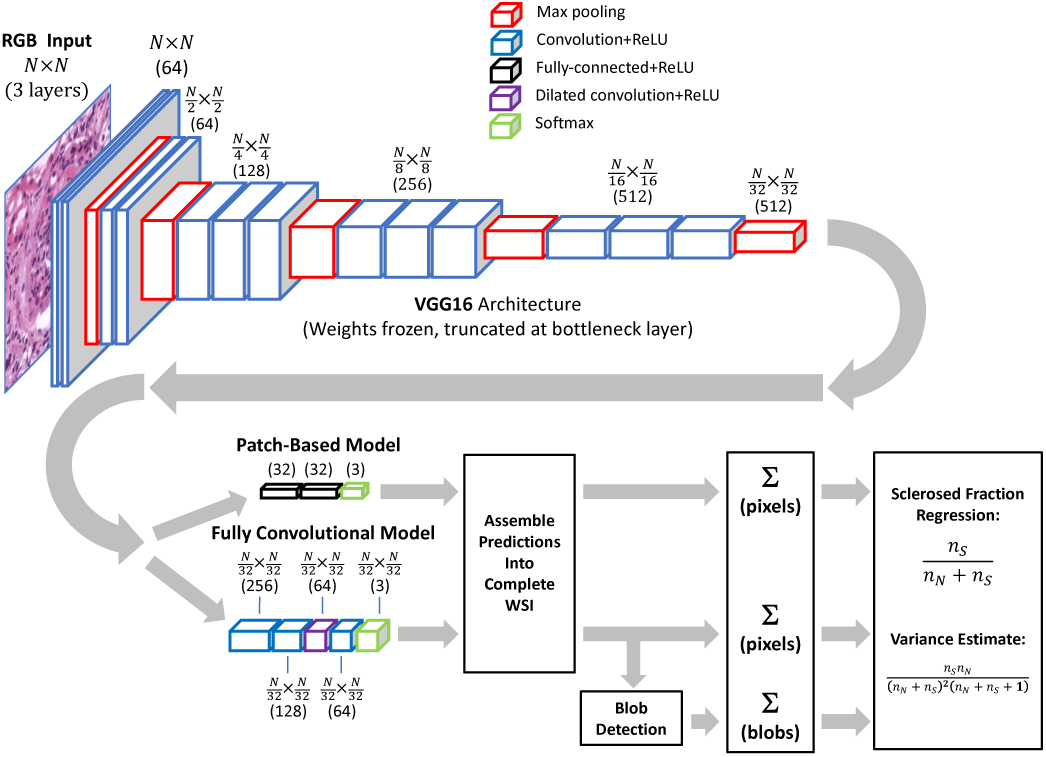
Data path used for computation of sclerosed glomeruli fraction. Both patch-based and fully-convolutional models utilize pretrained VGG16 architecture with frozen weights, truncated before bottleneck.

#### 1) Patch-Based Model

A patch-based CNN training approach was employed as a proof-of-concept to first demonstrate glomerulus differentiation in frozen H&E sections, but also to illustrate the pitfalls of applying this type of model to detect glomeruli in WSI. Image patches (448×448 pixels) centered on each labeled sclerotic and non-sclerotic glomerulus were cropped out of WSI for training. An additional 1932 randomly selected regions containing no glomeruli but at least a small fraction of non-whitespace were extracted for training on tubulointerstitial areas. Interstitial areas included tubules, vessels, inflammatory cells, and tubulointerstitium. The training set was augmented with random image flipping, 90°rotations, and small translations (0%—5% of image size). The pre-trained VGG16 CNN model [11] was adapted by removing the final fully-connected layers and replacing them with two 32-node fully-connected layers (with ReLU activation) and a 3-node classification layer with softmax activation; all weights in the lower convolutional layers were frozen and unmodified during training. The model was trained by minimizing the categorical cross-entropy loss using the Adam optimizer [43] with a batch size of 16 and a learning rate of 1*e*^−4^. The CNN was constructed using the Keras framework in Python and trained and tested using 6-fold cross-validation. Prior exploration using a test set indicated that stopping training at 5 epochs prevented overfitting while yielding satisfactory categorical accuracy, so this value was used in all cross-validation folds. The trained model from each fold was next applied to each of the associated WSIs withheld from training by sampling image patches from a window moved across the image in a raster pattern (448×448-pixels, 64-pixel stride), yielding a set of categorical probability values associated with each respective image patch. These values were assembled to yield a categorical probability map of the parent image downsampled by a factor of 64. Although it would have been preferable to use a 32-pixel stride in order to exactly match the output resolution of the fully convolutional model described below, the time required to generate the probability maps was prohibitive.

#### 2) Fully Convolutional Model

In addition to the patch-based model, we also trained a fully convolutional model based on VGG16 to label WSIs at higher resolution. Starting with the pre-trained VGG16 CNN with weights frozen below the bottleneck, we replaced the final fully-connected layers with two 1×1 convolutional layers (256 and 128 nodes, respectively), followed by a 64-node 3×3 dilated convolution layer [44], [45] (dilation rate=4) and another 64-node 5×5 convolutional layer. All convolutional layers used ReLU activation. Output was fed to a 3-node layer with softmax activation for classification into tubulo-interstitium, non-sclerosed glomerulus, and sclerosed glomerulus categories. Storing the activations of a fully convolutional network over an entire WSI is not feasible due to excessive memory requirements, therefore we adopted a sampling approach to training the model. For each training image, 1024×1024-pixel partially-overlapping image patches (stride=448) were extracted and presented to the model by weighted sampling. Classes were weighted in a ratio of 10:5:1 for sclerosed:non-sclerosed:tubulo-interstitial categories to approximately account for the relative incidence of pixel area represented by each class. The task of the CNN was to assign labels to individual pixels in the WSI to match the provided annotations. Because the model’s output was downsampled by a factor of 32 relative to the original image, the CNN was trained against a similarly downsampled annotation map (accomplished by nearest-neighbor sampling). The fully convolutional CNN was trained by minimizing the categorical cross entropy loss using the Adam optimizer with a batch size of 25 and learning rate of 1*e*^−4^ for 5 epochs. The model was trained and tested in a 6-fold cross-validation scheme. Prior exploration using a test set indicated that stopping training at 5 epochs prevented overfitting while yielding satisfactory categorical accuracy, so this value was used in all cross-validation folds. For each fold, the corresponding trained model was applied to the withheld set of WSIs by sampling image patches in a raster pattern from a sliding window (1024×1024-pixels) with a stride of 448 pixels, yielding a 32×32-pixel categorical probability map associated with each respective image patch. These patches were stitched together to yield a categorical probability map of the complete WSI, downsampled by a factor of 32.

## IV. Results

### A. Performance on Patches

The patch-based model’s performance in predicting image patch category was evaluated in terms of precision, recall, and F1-score averages over cross-validation runs. Results are shown in Table II. A normalized confusion matrix derived is shown in Table III. F1-score (the harmonic mean of precision and recall) for the non-sclerosed category is somewhat higher than for tubulo-interstium and sclerosed. This may be anticipated, given the greater variability in appearance associated with both the bulk tissue and the pathology (e.g., left panel of Figure 2). Examples of correctly and incorrectly identified glomeruli are shown in Figure 4. Note that the model correctly classifies non-sclerotic and sclerotic glomeruli even in the presence of preparation artifacts, staining variations, indeterminate Bowman’s space, and significant background whitespace within the image patch. While the patch-based model performed well in this proof-of-concept scenario, it is ultimately more important to gauge performance on WSI.

**Fig. 4:**
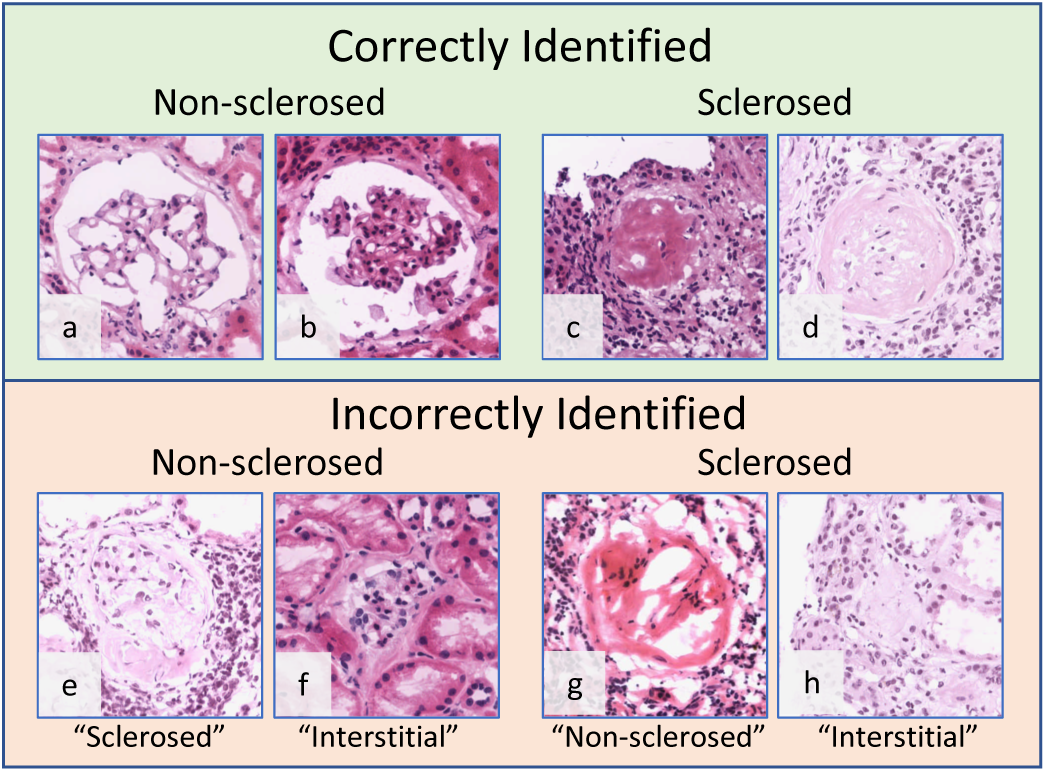
Examples of patch-based model predictions on image patches containing isolated glomeruli. Top: Highest scored correctly identified patches. The model correctly identified sclerosed and non-sclerosed glomeruli, even in the presence of variable stain intensity and glomerular appearance. Bottom: Lowest scored incorrectly labeled patches; predicted label is shown in quotes beneath each image.

**Table II.** Patch-based model performance when categorizing isolated glomeruli

|  | Precision | Recall | F1-score |
| --- | --- | --- | --- |
| Tubulointerstitial | 0.932 | 0.898 | 0.915 |
| Non-sclerosed | 0.932 | 0.962 | 0.947 |
| Sclerosed | 0.893 | 0.865 | 0.879 |

**Table III.** Normalized confusion matrix for patch-based model when categorizing isolated glomeruli

| Predicted\True | Tubulo-interstitial | Non-sclerosed | Sclerosed |
| --- | --- | --- | --- |
| Tubulointerstitial | 0.898 | 0.077 | 0.025 |
| Non-sclerosed | 0.024 | 0.962 | 0.014 |
| Sclerosed | 0.062 | 0.073 | 0.865 |

### B. Performance on WSI: Pixelwise Results

Selected WSIs are shown with their associated annotations and predicted probability maps for examples having large numbers of sclerosed glomeruli (Figure 5), small numbers of sclerosed glomeruli along with visible section folding artifact (Figure 6), and large regions of renal capsule (Figure 7), for both the patch-based and fully convolutional models. Probability magnitude is indicated by the brightness of color associated with each label (blue→non-sclerosed, red→sclerosed, tubulointerstitium not shown). The patch-based model does not generalize well to the task of segmenting glomeruli in WSI, especially in instances with small numbers of sclerosed glomeruli and prominent renal capsule. The fully convolutional model predictions, however, appear faithful in position and shape to the majority of annotated glomeruli. Additionally, the fully convolutional model’s glomerular labeling is much more focal in nature, whereas the patch-based model is often characterized by diffuse regions of positive labeling. It is also noteworthy that folding artifacts (see Figure 6a) have no apparent effect on the fully convolutional model’s performance.

**Fig. 5:**
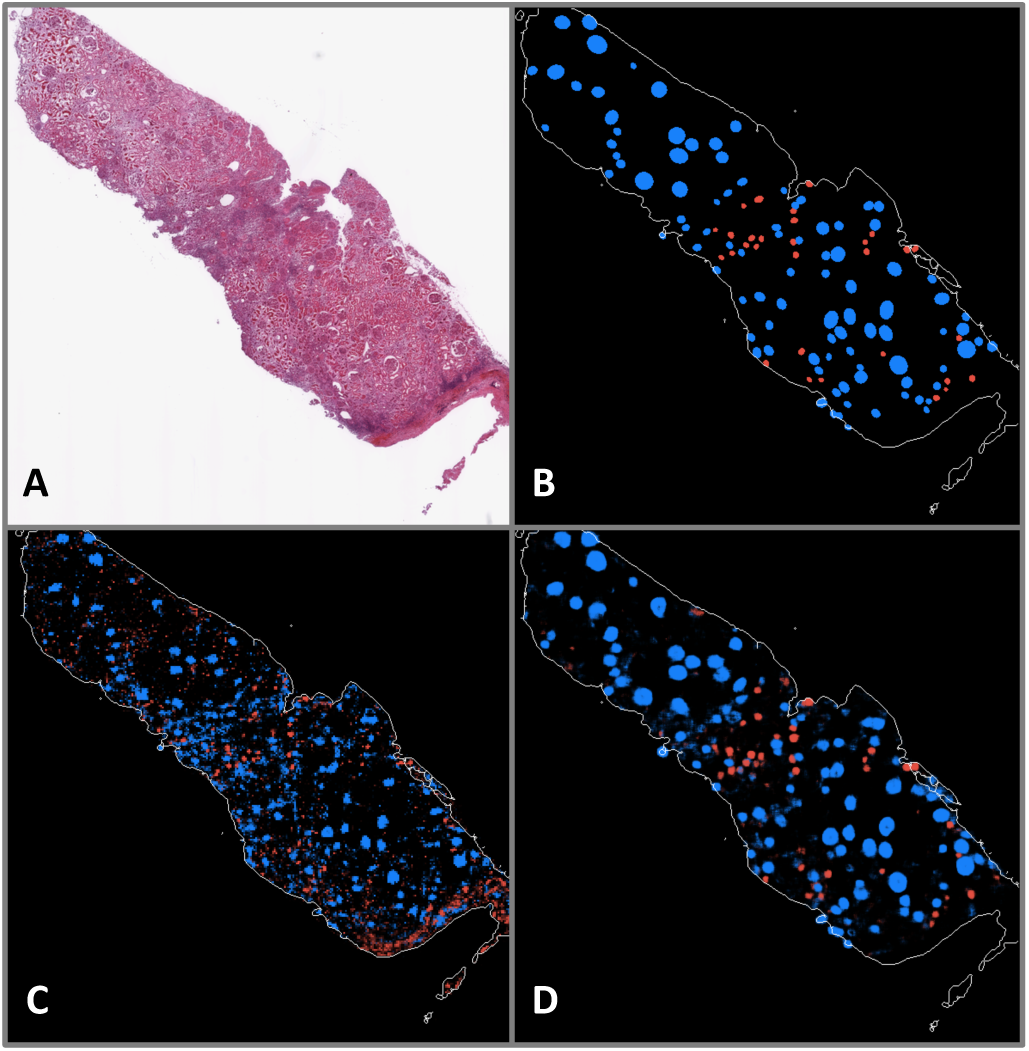
A: WSI exhibiting a large number of sclerosed glomeruli. B: Ground truth annotations indicating positions and shapes of non-sclerosed (blue) and sclerosed (red) glomeruli. C: Patch-based model prediction probability map. D: Fully convolutional model prediction probability map.

**Fig. 6:**
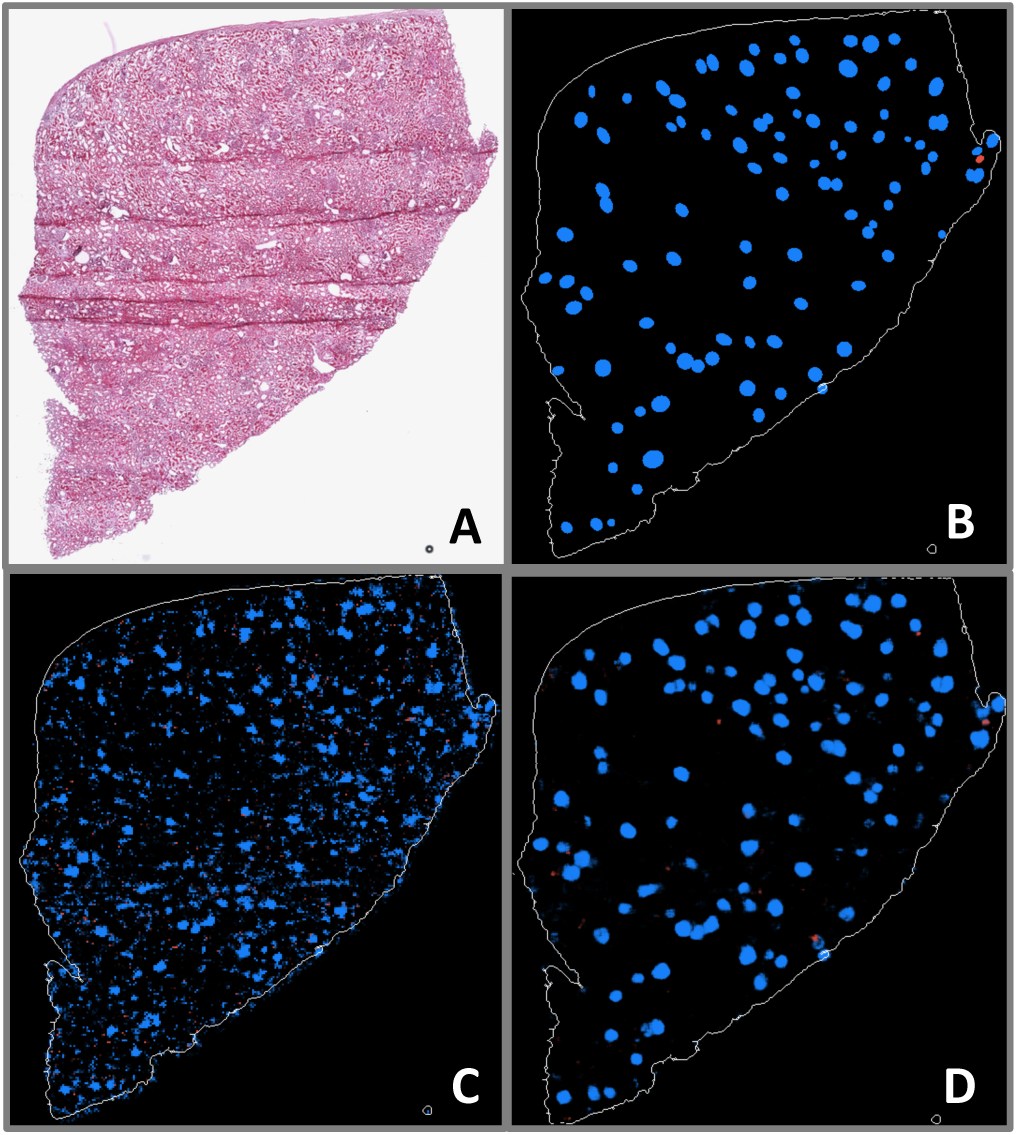
A: WSI exhibiting very few sclerosed glomeruli, as well as folding artifacts. B: Ground truth annotations indicating positions and shapes of nonsclerosed (blue) and sclerosed (red) glomeruli. C: Patch-based model prediction probability map. D: Fully convolutional model prediction probability map.

**Fig. 7:**
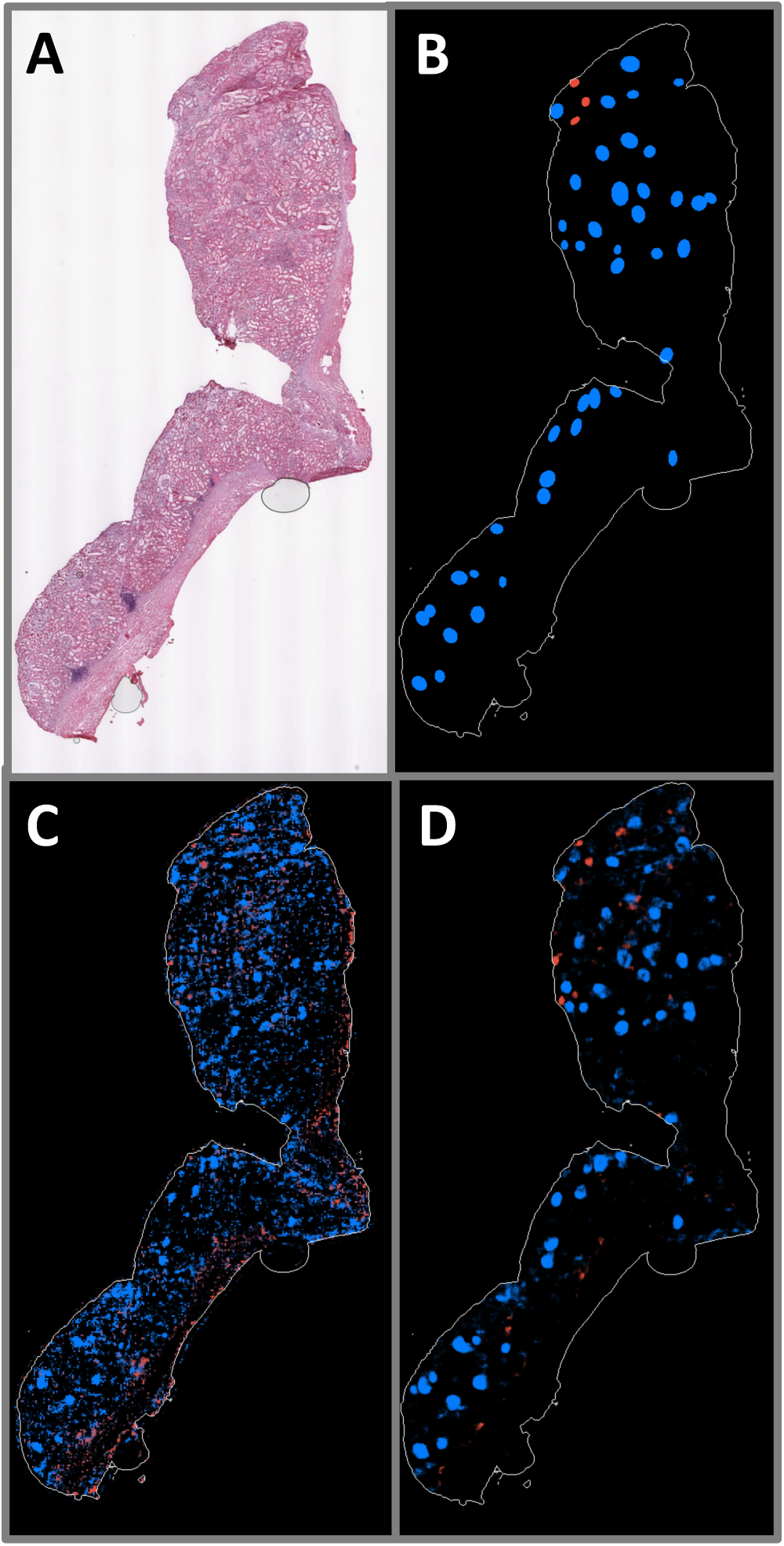
A: WSI exhibiting large region of renal capsule. B: Ground truth annotations indicating positions and shapes of non-sclerosed (blue) and sclerosed (red) glomeruli. C: Patch-based model prediction probability map. D: Fully convolutional model prediction probability map.

To quantify agreement between model predictions and pathologist annotations on a pixel-by-pixel basis, each pixel in the probability maps was assigned the categorical label associated with the highest probability at that point. Percent area fraction (Table IV) and intersection-over-union (IOU) (Table V) metrics were computed from the predicted label maps for all WSIs in each cross-validation fold. The IOU (also known as the Jaccard index) is computed by comparing the number of pixels in each category in which the predicted and annotated labels agree (intersection) divided by the total number of predicted and annotated pixels assigned a label for that category (union). These quantities were computed in aggregate for all pixels in the WSI predictions. Both models have high concurrence for tubulointerstitial pixels (Tables IV and V). Non-sclerosed areas were more reliably labeled by the fully convolutional compared to the patch-based model, as measured by area fraction. The fully convolutional model’s score was higher than the pathologists’ assessment by 28%, whereas the patch-based model overestimated by 58%. Model differences are most stark in the sclerosed category. The patch-based model drastically overestimates the sclerosed area (6.0x), compared to 1.7x overprediction for the fully convolutional model. IOU scores are even more telling, with the fully convolutional model having respectable scores of 0.59 and 0.36 for non-sclerosed and sclerosed areas, respectively, while the patch-based model returns IOU values of only 0.20 and 0.07, respectively.

**Table IV.**
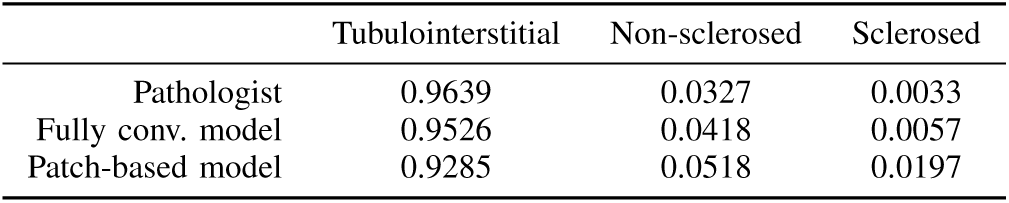
Pixel area fraction

**Table V.** IOU (by pixel) predicted by each model, referenced to pathologist annotations

|  | Tubulointerstitial | Non-sclerosed | Sclerosed |
| --- | --- | --- | --- |
| Fully conv. model | 0.9766 | 0.5949 | 0.3560 |
| Patch-based model | 0.9160 | 0.2017 | 0.0713 |

Kidney preimplantation biopsy evaluation requires the determination of the fraction of the number of sclerosed glomeruli in a WSI (given by *F* = *n*_*S*_/(*n*_*S*_+*n*_*N*_)). An equivalent, pixel-based surrogate measure was computed for comparison using the predicted label maps given by each model. The sclerosed fraction was computed as *F*_*pixel*_ = *n*_*S,pixel*_/(*n*_*S,pixel*_ + *n*_*N,pixel*_), where *n*_*S,pixel*_ and *n*_*N,pixel*_ are the number of pixels labeled as sclerosed and non-sclerosed, respectively. To assess the models’ accuracy in estimating the sclerosed fraction, we trained zero-intercept linear regression models. These linear models took *F*_*pixel*_ as input to be fit to *F*, as determined by pathologist annotations. The models were evaluated by *R*^2^ and root mean square error (RMSE) on the cross-validation test set.

The fully convolutional model showed greater correlation with percent global glomerulosclerosis (*R*^2^ = 0.828) compared with the patch-based model (*R*^2^ = −0.491). The mean slope of regression (averaged over cross-validation folds) for the patch-based model was 0.640, and 0.581 for the fully convolutional model. The resulting output is shown in Figure 8A. Error bars indicate the 95% confidence interval, assuming the sclerosed population fraction is characterized by a beta distribution. In practice, prior results have indicated poorer clinical outcome for donor kidneys with greater than 20% global glomerulosclerosis. Gray dotted lines are plotted at the 20% point on each axis, dividing the plot into quadrants. In this way, the plot can be read as showing that the model agrees with trained observers for samples lying in the lower left quadrant (organ more acceptable) and upper right quadrant (organ less acceptable). It can be clearly seen that the fully convolutional model output agrees with the pathologists’ assessments to a far greater degree than the patch-based model in this regard.

**Fig. 8:**
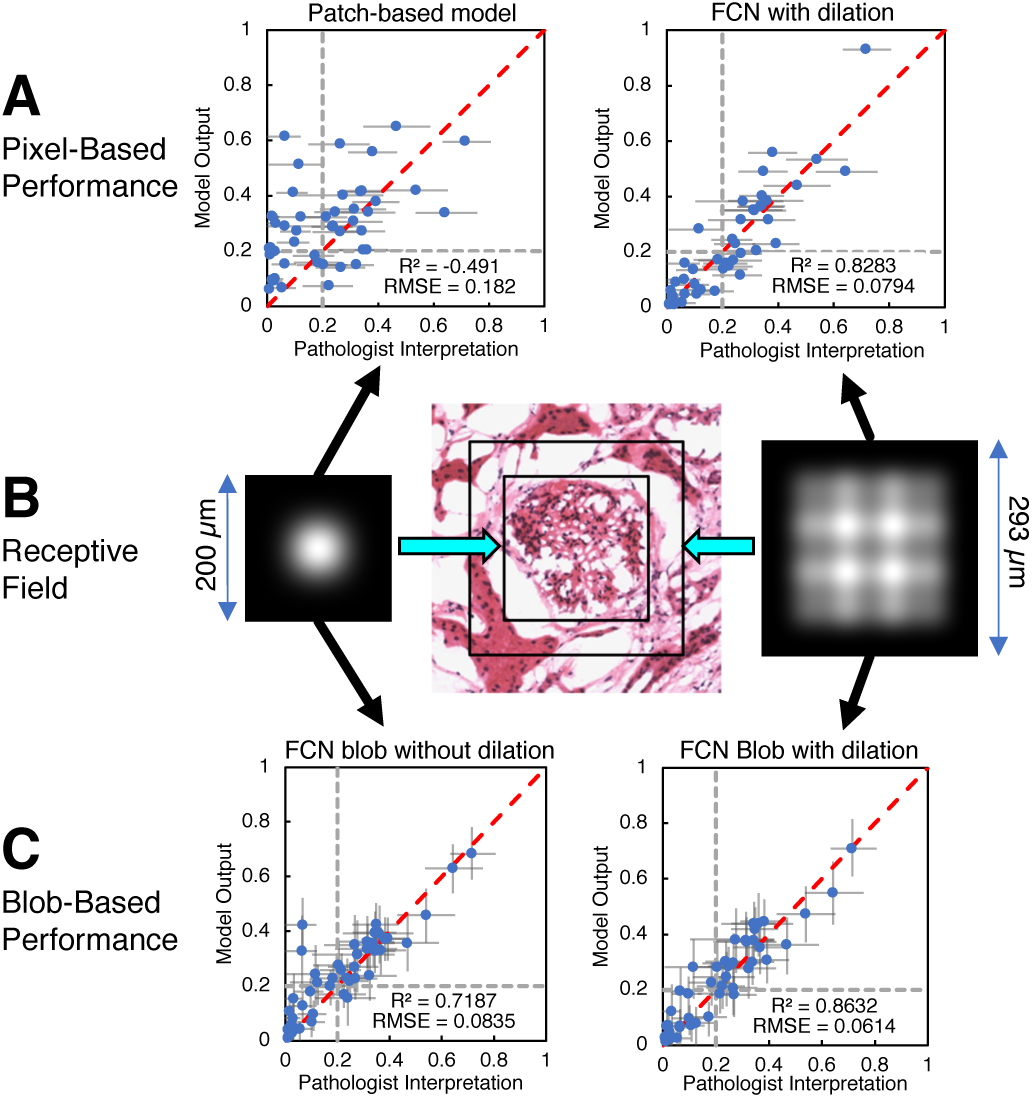
A: Pixel-wise model predictions of sclerosed glomerulus fraction vs. pathologists’ assessment for patch-based model (left) and fully convolutional model (right). Horizontal error bars indicate 95% confidence level for pathologist assessments. Dotted grey lines indicate a hypothetical clinical cutoff for rejection at 20% sclerosed. B: Receptive field intensity map for fully convolutionall model without dilated convolution layer (left) and with dilated convolution layer (right). Receptive field extent for both models are drawn to scale on image of normal glomerulus extracted from a WSI (center). C: Predictions of sclerosed glomerulus fraction vs pathologists assessment for fully convolutional model without dilated convolution layer (left) and with dilated convolution layer (right) after blob-detection postprocessing. Error bars indicate 95% confidence level, assuming the sclerosed and non-sclerosed glomeruli population is characterized by a beta distribution. Dotted grey lines indicate a hypothetical clinical cutoff for rejection at 20% sclerosed.

### C. Performance on WSI: Segmenting Glomeruli with Blob Detection Post-Processing

While the fully convolutional model’s pixelwise performance described above shows significant promise for evaluating slides in a global sense, identification of individual glomeruli is an important additional step for pathologists’ visual confirmation. In addition, identifying individual glomeruli enables the model to generate position and shape information for input to other glomerulus image characterization procedures. Only the fully convolutional model was utilized in this procedure, because of its superior performance and resolution relative to the patch-based model. A conventional Laplacian-of-Gaussian (LoG) blob-detection algorithm [46] was used to process the fully convolutional model’s probability map predictions for identification of the locations of sclerosed and non-sclerosed glomeruli. The LoG algorithm (implemented in the scikit-image Python library [47]) outputs position and approximate radius of detected objects. Examples of output from blob detection are shown in Figure 9 for the previously depicted WSIs in Figures 5-7. In the panels showing glomerulus detections, solid circles indicate confirmed matches to annotations, X’s mark incorrect glomerulus detections, and open rectangles indicate annotated glomeruli that were overlooked by the model.

**Fig. 9:**
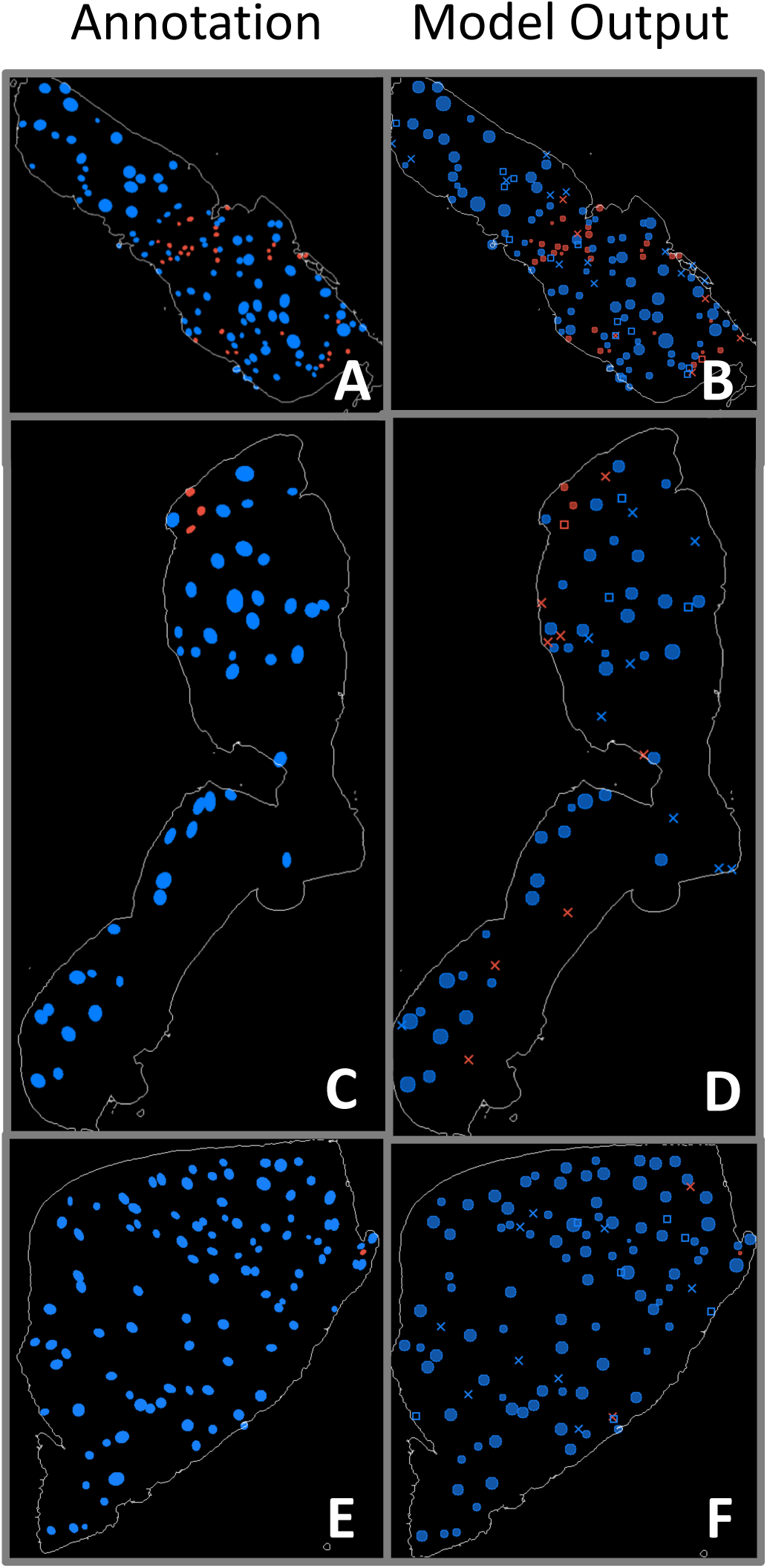
Panels A, C, E: WSI annotations indicating positions and shapes of non-sclerosed (blue) and sclerosed (red) glomeruli. Panels B, D, F: Corresponding blob detection results using fully convolutional model prediction label maps as input. Solid circles indicate confirmed matches with annotations, X’s mark incorrect glomerulus detections, and open rectangles indicate annotated glomeruli that were overlooked by the model.

Accuracy metrics for the predictions are given in Table VI. A detected blob was considered positively identified if its center was located within the area of a region in the annotation map, and if it had the same label as that annotated region. The total number of detected sclerosed and non-sclerosed glomeruli were modestly higher than the pathologists’ assessments, differing by 14.9% and 8.9%, respectively. Precision and recall were higher for non-sclerosed glomeruli than for sclerosed, reflected in F1 scores of 0.8475 and 0.6492, respectively. Linear regression of the sclerosed and non-sclerosed glomeruli were modestly higher than the pathologists’ assessments, differing by 14.9% and 8.9%, respectively. Precision and recall were higher for non-sclerosed glomeruli than for sclerosed, reflected in F1 scores of 0.8475 and 0.6492, respectively. Linear regression of the sclerosed fraction *F*_*blob*_ = *n*_*S,blob*_/(*n*_*S,blob*_ + *n*_*N,blob*_) (where *n*_*S,blob*_ and *n*_*N,blob*_ are number of sclerosed and non-sclerosed glomeruli obtained from the blob-detected probability maps) versus the pathologists’ assessment for each WSI in the training set was computed and used to regress the blob-detected probability maps in the cross-validation test set (Figure 8C, right). The mean coefficient of regression (averaged over cross-validation folds) was 0.978. Both *R*^2^ and RMSE were improved versus the equivalent pixel-based metrics for this model (*R*^2^: 0.863 vs. 0.828, RMSE: 0.061 vs. 0.079). Note that the fully convolutional model’s RMSE value approached the intrinsic error of the pathologists assessment (0.043), computed from the square root of the mean value of sclerosed fraction variance.

**Table VI.** Accuracy metrics computed from the fully convolutional model’s probability maps of wsis after blob detection.

|  | Non-sclerosed | Sclerosed |
| --- | --- | --- |
| Pathologists' count | 2997 | 870 |
| Predicted count | 3264 | 1000 |
| Correct (true positive) | 2653 | 607 |
| Incorrect (false positive) | 611 | 393 |
| Missed (false negative) | 344 | 263 |
| Precision | 0.8128 | 0.6070 |
| Recall | 0.8852 | 0.6977 |
| F1-score | 0.8475 | 0.6492 |

### D. Misclassified Glomeruli

A list of misclassified glomeruli images from the test set were submitted to the pathologists responsible for the ground truth annotations in order to determine the rate of actual misclassified glomeruli, and also to gauge the incidence of glomeruli that may have escaped initial inspection. Of the false positive predictions represented in Table VI, 34 glomeruli were determined to have been overlooked in the annotation process, 7 were judged to be misidentified by the pathologists, and 13 were correctly identified by the fully convolutional model but positioned immediately adjacent to (rather than atop) the annotated glomeruli. Of the false positive predictions that were not tubulointerstitium, 199 were areas of cyst or tattered regions associated with frozen artifact, 83 were vessels, 6 were areas of thick or folded tissue, 3 were instances of hyaline or atrophied tubules, 7 were borderline cases not clearly normal or sclerosed, and 13 were areas of where the detected blob encompassed multiple glomeruli.

## V. Discussion

Several factors were key to the success of the fully convolutional model relative to patch-based model. Because the fully convolutional network’s output maintained pixel-wise fidelity to the downsampled annotation map, network training and prediction occurred at a higher resolution than the patch-based approach. In particular, the fully convolutional model was exposed to a far greater variety of tissue configurations, in which all areas of the input image patches were able to inform the model during training.

The use of a dilated convolutional layer [44], [45] also increased the receptive field of the network. In order to illustrate the effect of the dilated convolution, we constructed and trained a model that replicated the same structure as the fully convolutional model, but replaced the dilated convolution layer with a normal convolution layer. Additionally, we implemented linearized versions of both fully convolutional models with all weights set to unity, biases set to zero, max pooling layers replaced with average pooling, and activations set to the identity function. An “image” array initialized to zero, except for a single point set to a constant nonzero RGB value, was used as input to the linearized models. The central value of each linearized model’s output array was stored in a result array at the same position as the nonzero point in the input array. Linearized model outputs were computed for each realization of the input as the nonzero point was moved over every position in the array. The extent of the receptive field was determined by the nonzero points in the result array. The results are shown in Figure 8B, and can be interpreted as idealized “point spread functions” that map the relative contribution of nearby input pixels to an individual output point (ignoring learned weights in this illustration). Regression plots for sclerosed glomerulus fraction using blob detection post-processing for both models show the improvement in prediction agreement with pathologist assessments enabled by the dilated convolution layer (Figure 8C). The two-tailed p-value for the difference between the correlation coefficients of the two models was 0.056, approaching significance at the 95% level. The difference in prediction between the models was most evident for samples with large capsule areas, in which the model without the dilated convolution layer erroneously inferred sclerosed glomeruli adjacent to fibrous regions. We speculate that the model utilizing dilated convolution was better able to infer regional context because of its larger receptive field.

Although the size of the final model’s receptive field was increased, we found that, for a given input image patch, the central area of the model prediction output array typically exhibited higher correlation with the corresponding annotation region than areas closer to the output array’s perimeter. Thus, the stride used in selecting image patches for training and prediction was chosen to be 448 pixels so as to enable sample overlap of 576 pixels (56% overlap for 1024-pixel square arrays). Only the central, non-overlapping portions of the prediction patches were used to assemble the final probability maps, which markedly improved results when compared to the use of, for instance, an 896-pixel stride.

Of the glomeruli misidentified by the fully convolutional model that were not associated with generic tubulointerstitium, by far the largest number occurred in areas centered on cysts, vessels, or holes. Many of these regions included somewhat circular features similar in scale to glomeruli, which may have led to the misclassification. It is possible that the model’s performance could be further enhanced by labeling and training on WSIs annotated with these categories. Very few instances of misclassification occurred because of tissue folding, however, suggesting that the model was insensitive to this artifact.

Using blob detection to infer glomerulus presence from the pixel map incurs the same potential drawbacks as those found in other morphological detection methods, namely the need to determine appropriate input parameters based on domain knowledge. In this case, parameters for minimum and maximum radius and minimum intensity threshold were fixed at levels which yielded reasonable discrimination between closely spaced glomeruli, and which excluded low-intensity regions or objects too small to be glomeruli. In making these choices, however, it is possible that additional errors may arise that can obscure the quality of the underlying data. Figure 10 highlights one type of error in which adjacent predicted glomerular regions are not resolved by the LoG algorithm, leading to a false negative result for one of the adjoining glomeruli in spite of qualitatively correct labeling displayed by the fully convolutional model’s probability map. Nevertheless, this post-processing step allows glomerulus-level estimation of the model’s fitness for slide evaluation through direct comparison with the pathologists’ estimates of glomerulus populations. We anticipate that future work will include means of accounting for and training to glomerulus population in the network architecture.

**Fig. 10:**
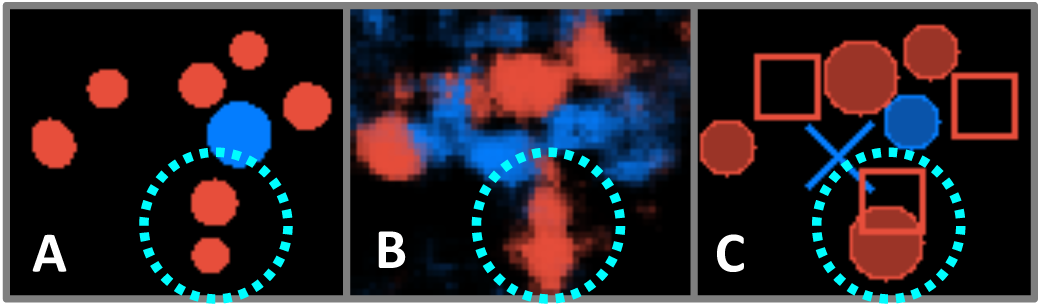
An example blob detection error highlighted within dotted cyan circle. A: Image patch extracted from WSI annotation map, indicating positions and shapes of non-sclerosed (blue) and sclerosed (red) glomeruli. B: Probability map for the same image patch. C: Blob detection results using fully convolutional model prediction label map as input. Solid circles indicate confirmed matches with annotations, X’s mark incorrect glomerulus detections, and open rectangles indicate annotated glomeruli that were overlooked by the model. The blob detection algorithm fails to differentiate the model’s correctly-identified adjoining sclerosed glomeruli in this instance.

In the long run, we aim to measure the extent to which this model reduces the inter-observer variability of pathologist assessments, and the intrinsic error associated with assessing percent globular sclerosis off of a single WSI. Though very preliminary, it appears that the variability of the model’s output is lower across technical replicates (Table VII). Two of the kidneys associated with this study had five or more associated WSIs, and were, therefore, well-controlled technical replicates. In both series, the average value of the model prediction was in close agreement with the pathologists’ determination; moreover, the standard deviation of the model output was lower than for the pathologist assessment, suggesting that the model may be able to decrease evaluation variability.

**Table VII.** Global glomerulosclerosis estimates for kidneys with well-controlled technical replicates (5 or more sections).

|  | Pathologist |  | Model |  |
| --- | --- | --- | --- | --- |
|  | mean | SD | mean | SD |
| Case 1 | 24.2 | 5.2 | 25.5 | 4.8 |
| Case 2 | 2.6 | 1.7 | 2.7 | 0.3 |

## VI. Conclusion

Two deep-learning models were adapted from a pretrained CNN for the purpose of glomerular segmentation and classification of glomeruli in frozen-section whole-slide images. This task is critical for the time-sensitive evaluation of donor kidneys before transplantation. The initial patch-based architecture was able to robustly categorize sclerosed glomerular image patches from non-sclerosed after training with a relatively small set of samples, showing the utility of transfer learning with a general-purpose image classification CNN. Applied to whole-slide images, the patch-based model was outperformed by a fully convolutional model based on the same pretrained CNN. A blob detection post-processing step was used to generate discrete maps of glomeruli and their associated class (sclerosed or non-sclerosed). Percent global glomerulosclerosis, a key metric used in grading kidneys for transplant suitability, indicated performance for the fully convolutional CNN nearly equivalent to that of a board-certified clinical pathologist. We are optimistic that the methodology described here has the potential to be an essential part of the workflow for transplant evaluation in the clinical setting.

## Acknowledgment

The authors would like to thank the Institute for Informatics at Washington University in St. Louis, the Mid-America Transplant Foundation, and the National Institutes of Health for their support of this work. Contributions: J.N.M wrote the manuscript and, with M.K.M, developed the software used in the processing pipeline; M.K.M developed final architecture of the fully convolutional model; S.K. and J.P.G. annotated all WSIs and assessed model results; T.-C.L. and T.S.S provided additional guidance in clinical- and pathology-related matters; S.J.S. directed the project and provided guidance for deep learning tasks; J.P.G., T.-C.L. and S.J.S. obtained funding.

